# The nonlinearity of regulation in biological networks

**DOI:** 10.1101/2021.12.22.473903

**Authors:** Santosh Manicka, Kathleen Johnson, Michael Levin, David Murrugarra

## Abstract

The extent to which the components of a biological system are (non)linearly regulated determines how amenable they are to therapy and control. To better understand this property termed ‘regulatory nonlinearity’, we analyzed a suite of 137 published Boolean network models, containing a variety of complex nonlinear regulatory interactions, using a probabilistic generalization of Boolean logic that George Boole himself had proposed. Leveraging the continuous-nature of this formulation, we used Taylor decomposition to approximate the models with various levels of regulatory nonlinearity. A comparison of the resulting series of approximations of the biological models with appropriate random ensembles revealed that biological regulation tends to be less nonlinear than expected, meaning that higher-order interactions among the regulatory inputs tend to be less pronounced. A further categorical analysis of the biological models revealed that the regulatory nonlinearity of cancer and disease networks could not only be sometimes higher than expected but also relatively more variable. We show that this variation is caused by differences in the apportioning of information among the various orders of regulatory nonlinearity. Our results suggest that there may have been a weak but discernible selection pressure for biological systems to evolve linear regulation on average, but for certain systems such as cancer, on the other hand, to also evolve more nonlinear rules.

## 1 Introduction

How nonlinear is the regulation of the components of biological networks? That is, to what extent do the biochemical components of these networks non-independently interact (Fig 1) in influencing downstream processes and network behavior. Research on the ‘nonlinearity’ front has hitherto focused on its various manifestations in the dynamics of biological systems, such as chaos, bifurcation, multistability, synchronization, patterning, dissipation, etc.[1], but a characterization of *regulatory* nonlinearity among the components of the underlying systems that give rise to those phenomena is lacking. A more complete understanding of biological regulatory nonlinearity would not only yield insights into their design principles [2] but also have theoretical implications ranging from canalization to control [3, 4] and practical implications for biomedical therapy, synthetic biology, etc. [1, 5]. A good example of this concerns the mapping between molecular or genetic information and the resulting system-level anatomical structure and function of an organism. Advances in regenerative medicine and synthetic morphology require rational control of physiological and anatomical outcomes [6], but progress in genetics and molecular biology produce methods and knowledge targeting the lowest-level cellular hardware. There is no one-to-one mapping from genetic information to tissue- and organ-level structure; similarly, ion channels open and close post-translationally, driving physiological dynamics that are not readily inferred from proteomic or transcriptomic data. System-level properties in biology are often highly emergent, with gene-regulatory or bioelectric circuit dynamics connecting initial state information and transition rules to large-scale structure and function. Thus, the difficult inverse problem [7] of inferring outcomes and desirable interventions across scales of biology illustrates some of the fundamental questions about the directness or nonlinearity of encodings of information, as well as the importance of this question for practical advances in biomedicine and bioengineering that exploit the plasticity and robustness of cellular collectives. Many deep questions remain about the potential limitations and best strategies to bridge scales for prediction and control in developmental, evolutionary, and cell biology. To that end, we introduce here a formal characterization of the nonlinearity of models of biological regulatory networks, such as those often used to describe relationships between regulatory genes. Specifically, we consider a class of discrete models of biological regulatory systems called “Boolean models” that are known for their relative simplicity and tractability compared to continuous ordinary differential equation-based (ODE) models [8].

**Figure 1:**
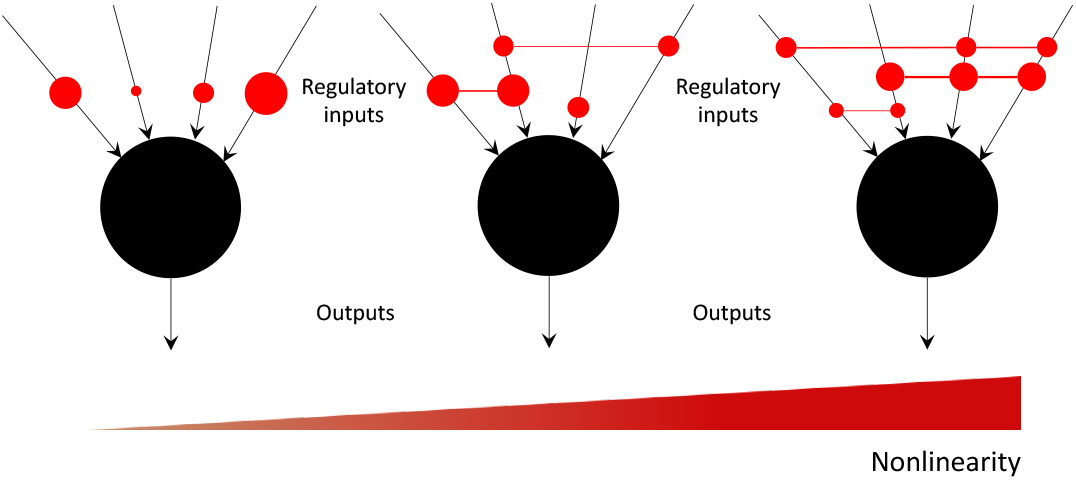
An illustration of the concept of regulatory nonlinearity. Each black circle represents a generic biochemical component such as a gene, transcription factor, enzyme, etc., regulated by a set of inputs (also biochemical components) and generates a generic output such as concentration level, strength, etc. Non-zero interactions among the inputs are represented by red circles connected by red lines, with the total number of possible interactions for a node with *k* inputs equal to 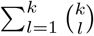. The size of the red circles and the width of the connecting lines represents the weight of the interactions. Independent inputs are represented by unconnected red circles. The degree of nonlinearity would thus be expected to increase, though not necessarily linearly, from left to right, as the numbers and the strengths of the regulatory interactions increase. One could also visualize these local interactions in a broader network-context as “hypergraphs” [17].

A Boolean network is a discrete network model characterized by the following features. Each node in a Boolean network can only be in one of two states, ON or OFF, which represents the expression or activity of that node. The state of a node depends on the states of other input nodes which are represented as a Boolean rule of these input nodes. Many of the available Boolean network models were created via literature search of the regulatory mechanisms and subsequently validated via experiments [9]. Some of the publicly available models were generated via network inference methods from time course data [3].

Previous studies have found that certain characteristic features of the biological Boolean models, such as the mean in-degree, output bias, sensitivity and canalization, tend to assume an optimal range of values that support optimal function [10, 11]. Here we study a new but generic feature of complex systems termed “regulatory nonlinearity” that we broadly define as the degree to which the inputs to the components of a complex interact. To characterize the regulatory nonlinearity of Boolean networks we formalize an approach to generalizing Boolean logic by casting it as a form of probability, which was originally proposed by George Boole himself [12]. We leverage the continuous nature of these polynomials to decompose a Boolean function using Taylor-series and reveal the distinct layers of its regulatory nonlinearity (Fig 2). Various other methods, both discrete and continuous, of decomposing Boolean functions exist, such as Reed-Muller, Walsh spectrum, Fourier, discrete Taylor and fuzzy logic [13, 14, 15, 16]. Our continuous Taylor decomposition method is distinct in that it offers a clear and systematic way to characterize nonlinearity.

**Figure 2:**
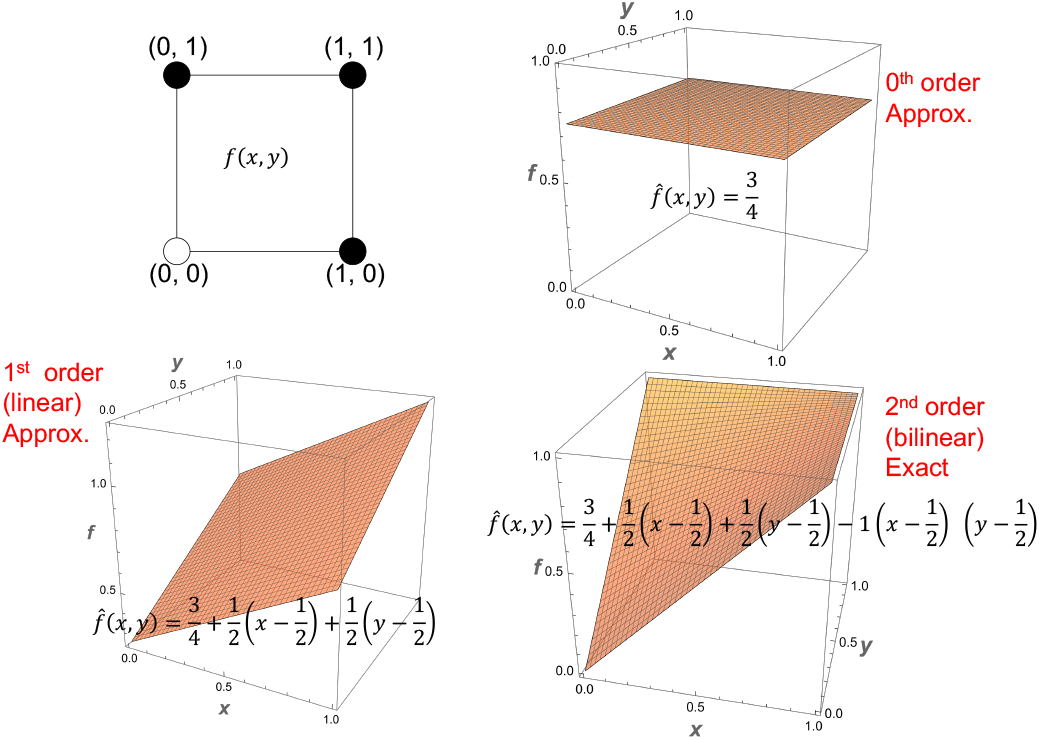
The various approximations of a Boolean function in increasing order of nonlinearity. The logical OR function is represented as a 2D hypercube (top left) with the coordinate values representing input combinations and the color of the circles representing the corresponding outputs (white=0, black=1) and is approximated using Taylor decomposition as the 0^*th*^ order approximation (top right) showing only the first term, the mean output bias; the 1^*st*^ order approximation (bottom left) including the linear terms; and finally the 2^*nd*^ order exact form (bottom right) including all the terms.

Is the regulatory nonlinearity of biological systems special? To that end we specifically ask: 1) how well could biological Boolean models be approximated, that is, faithfully represented with only partial information containing lower levels of nonlinearity relative to that of the original?; 2) is there an optimal level of regulatory nonlinearity, characterized by maximum approximability, that these models may have been selected for by evolution?; and 3) do different classes of biological networks show characteristically different optimal levels of regulatory nonlinearity? To answer these questions, we first approximate the biological models by systematically composing the various nonlinear layers resulting in a sequence of model-approximations with increasing levels of nonlinearity. We then estimate the accuracy of these approximations by comparing the outputs of their simulations with that of the original unapproximated model. We then construct an appropriate random ensemble for each biological model and compare their mean accuracies for fixed levels of approximation. The main idea is that a biological model that is more approximable than expected for a particular level of nonlinearity would mean that the network may have been optimized for that level nonlinearity. Finally, we classify the biological networks into various categories and compare their approximabilities to identify any category-dependent effects.

## Methods

### Probabilistic generalization of Boolean logic

Here we provide a continuous-variable formulation of a Boolean function by casting Boolean values as probabilities, thus transforming it into a pseudo-Boolean function [18]. Consider random variables *X*_*i*_ : {0, 1} → [0, 1], *i* = 1, …, *n*, with Bernoulli distributions. That is, *p*_*i*_ = *Pr*(*X*_*i*_ = 1) = 1 − *Pr*(*X*_*i*_ = 0) = 1 − *q*_*i*_, for *i* = 1, …, *n*. Let *X* = *X*_1_ × · × *X*_*n*_ be the product of random variables and *f* : *X* → {0, 1} a Boolean function. Let 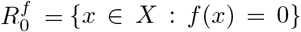 and 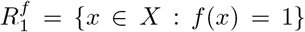. Note that *X* is a disjoint union of 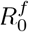 and 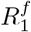. Then, 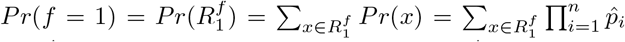 where 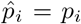 and 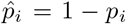.

Let 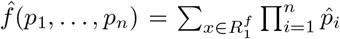. Thus, 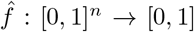 is a continuous-variable function. The following theorem shows that 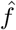 is a generalization of *f* in the sense that 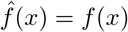 for all *x* ∈ { 0, 1} ^*n*^; proof is provided in SI Appendix.

#### Theorem 1.1.

*For discrete values of x*_*i*_ ∈ {0, 1}, *i* = 1, …, *n, we have* 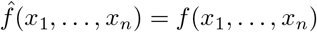.

#### Corollary 1.2.

*If p*_*i*_ = 1*/*2 *for all i* = 1, …, *n, then* 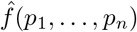 *is the output bias of f*.

#### Example 1.3.

*Consider the AND, OR, XOR, and NOT Boolean functions given in Table 1*. *The continuous-variable generalization of f*_1_, *f*_2_, *f*_3_, *and f*_4_ *are:* 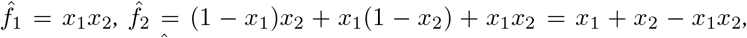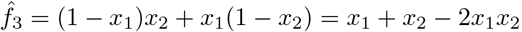, *and* 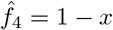.

**Table 1:**
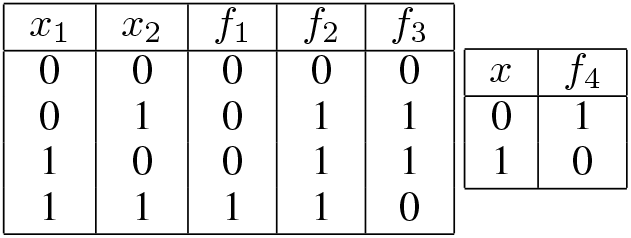
Truth tables of basic Boolean functions.

Note that the above expressions have previously been derived via other (not probability-based) means [15, 16].

### Taylor Decomposition of Boolean functions

Since 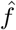 is a continuous-variable function, we can calculate its Taylor expansion. And since 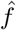 is a square-free polynomial, its Taylor expansion is finite and simplified (any term containing multiple derivatives of the same variable is zeroed out), as described in Proposition 1.4 using the standard multi-index notation. Let *α* = (*α*_1_, …, *α*_*n*_) where *α*_*i*_ ∈ {0, 1}.

We define 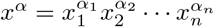, and 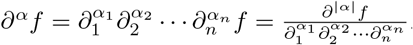.

#### Proposition 1.4.

*For p* ∈ [0, 1]^*n*^, *we have*

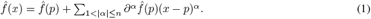

Note that 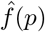 (*p*) in Equation 1 is the output bias of *f* as was seen in Corollary 1.2. A natural choice for *p* is *p* = (1*/*2, …, 1*/*2) as it represents an unbiased selection for each variable and it also gives the output bias of the function. Such unbiased choices are not available for the discrete case. Our continuous formulation thus offers such unique advantages over the discrete Taylor decomposition, as it’s a natural generalization of the latter. The Taylor decomposition can be used to approximate a Boolean function by considering a subset of the terms. For example, a linear approximation consists of terms only up to |*α*| ≤ 1, a bilinear approximation up to |*α*| ≤ 2, etc., up until |*α*| ≤ *n* where it ceases to be an approximation and provides an exact decomposition of 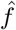. A visual illustration is provided in Figure 2. The approximation order of a Boolean *network* could therefore vary between its minimum and maximum in-degrees (number of inputs per node).

#### Example 1.5.

*Consider the continuous generalizations of the AND, OR, XOR and NOT functions given in Example 1*.*3 The corresponding Taylor expansions using Equation 1 and using the derivatives shown in Table 2 with p* = (1*/*2, 1*/*2) *are:* 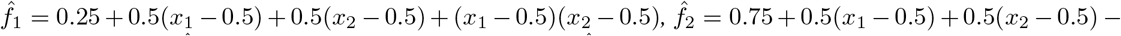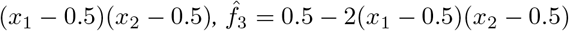, *and* 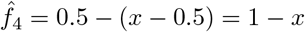.

**Table 2:**
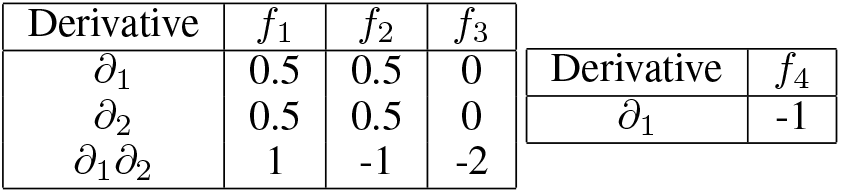
Values of partial the derivatives in the Taylor decompositions of the generalizations of basic Boolean functions.

| Derivative | $f_1$ | $f_2$ | $f_3$ | Derivative | $f_4$ |
| --- | --- | --- | --- | --- | --- |
| $\partial_1$ | 0.5 | 0.5 | 0 | $\partial_1$ | -1 |
| $\partial_2$ | 0.5 | 0.5 | 0 | | |
| $\partial_1 \partial_2$ | 1 | -1 | -2 | | |

Note that 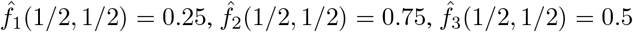, and 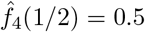 in the above equations are the output biases of the AND, OR, XOR, and NOT functions respectively. Also note that both the AND and OR functions contain the linear and the second order terms in their Taylor decomposition while the XOR function only contains the second order term. This difference is because both the AND and OR functions are monotone while XOR is not since it requires both inputs to be known.

### Approximability of a model

We considered a suite of Boolean network models of biochemical regulation from two sources, namely the *cell collective* [9] and reference [3]. This suite consists of 137 networks with the number of nodes ranging from 5 to 321. The mean in-degree of these models ranges from 1.1818 to 4.9375 with the variances ranging between 0.1636 and 9.2941, while the mean output bias is limited to the range [0.1625,0.65625] with the variances between 0.0070 and 0.0933. For each biological model an ensemble of 100 randomized models was generated each of whose connectivity and output biases were set to be the same as the former, with only the logic rules randomized. This set, referred to as the ‘constrained ensemble’, facilitates benchmarking the role of regulatory nonlinearity in the biological rules. To accurately assess the difference between the biological models and the constrained ensembles, we considered a baseline set known as the ‘unconstrained ensemble’ relative to which those differences were computed. This ensemble preserved neither the connectivity nor the output bias but had them bootstrap-sampled from the corresponding distributions characterizing the associated biological models. Taylor decomposition was then applied to both the biological models and the random ensembles and all possible nonlinear approximations were computed by considering terms starting from the linear order accumulating up to the maximum possible nonlinear order. Both the biological models and the random ensembles were then simulated using a set of 1000 randomly chosen initial states iterated through 500 update steps for all orders of approximation; the same initial conditions were used for a given biological model and the random ensemble. The states of the variables were restricted to the interval [0,1] at every step during the simulations by simply resetting them to the nearest boundary of the interval whenever they crossed it. The mean approximation error (MAE) of each model is defined as the mean squared error between the exact Boolean states and the approximated probabilistic states at the end of the simulations; for the random ensembles a single average MAE was computed. It varies between 0.0 and 1.0. The “percentage change in MAE” (PMAE) of the biological models and the constrained ensembles is defined as the percentage difference between the respective MAE and the corresponding unconstrained MAE; it can vary between -100.0 and positive infinity. The “approximability” of a biological model is represented by the difference between the PMAE of the corresponding constrained ensemble and that of itself; it can vary between negative infinity and positive infinity. Hence, the more negative the PMAE is for biological models compared to that of constrained ensembles the more approximable they are deemed to be and the more unique their regulatory nonlinearity is. An illustration of how approximability is computed is provided in Figure 3.

**Figure 3:**
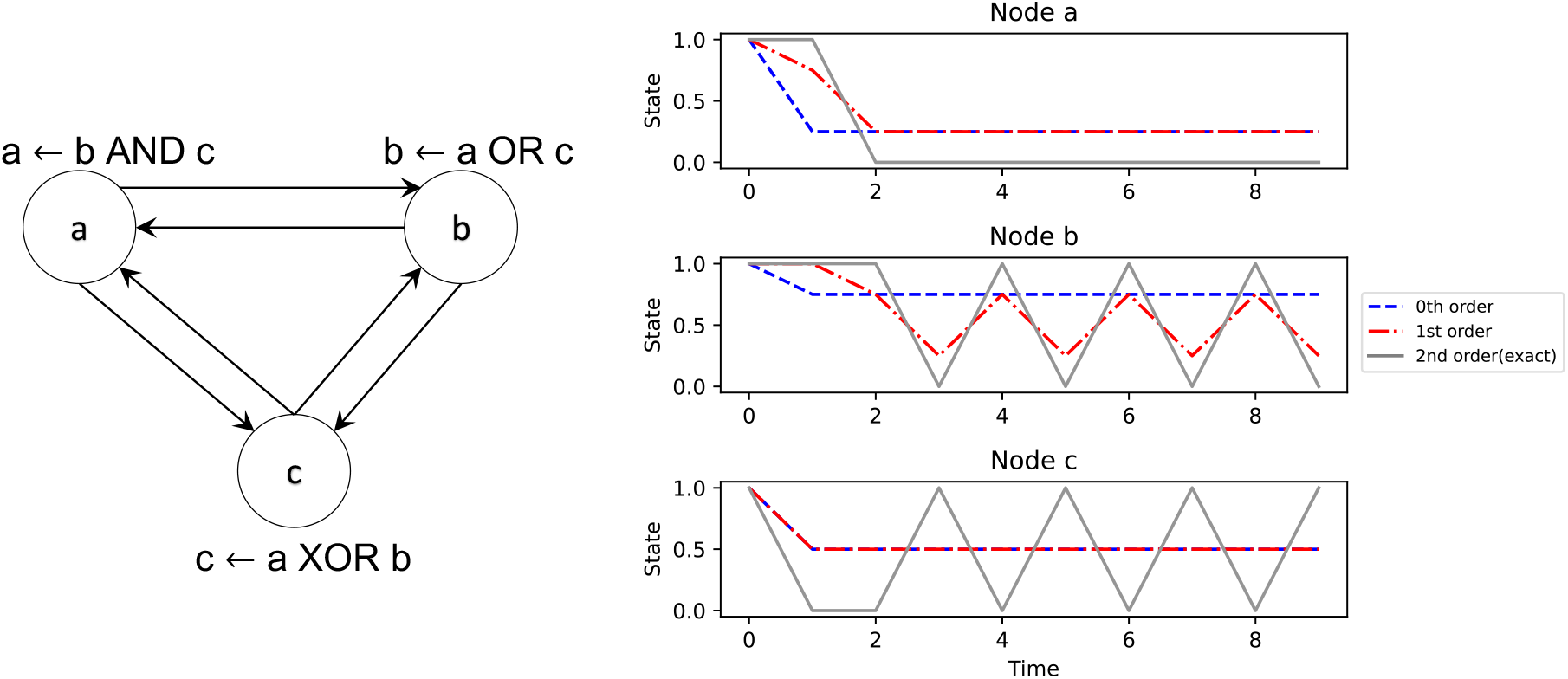
An illustration of the relationship between nonlinearity and approximability using a simple 3-node Boolean network. It shows that the higher the order of nonlinearity of the network the more approximable it is with respect to the exact dynamics. Left: An example 3-node network utilizing all three 2-input Boolean functions. Right: The network is simulated from a single initial state of (1,1,1). Here the dynamics of the 0^*th*^ order network (blue) least matches the exact Boolean trajectory (grey), while 1^*st*^ order network (red) is a slightly better match. The 2^*nd*^ order network (grey) is exact since it includes all the Taylor terms up to the maximum possible order of 2 (the maximum in-degree of any node). The closer the approximated average dynamics to the exact dynamics, the lower the MAE and the higher the approximability of the former. Notice also that while for node ‘a’ the 0^*th*^ order network is almost as good as the 1^*st*^ order network, it’s the opposite for node ‘c’ in that its 1^*st*^ order approximation is as poor as the 0^*th*^ order; this is because ‘c’ implements the XOR function whose 1^*st*^ order derivatives are zero.

### Classification of biological models

To identify any differences among the approximabilities of different types of biological networks we sought to classify them. Since there are multiple ways to classify biological networks, we chose two classifications so that: 1) they are as orthogonal as possible to each other; and 2) each classification has an appropriate number of (neither too few nor too many) categories. Classification 1 (C1) follows the “pathway ontology” (PW) [19] where the networks are grouped into five categories (Figure 5(a)), namely biochemical (*n* = 13), signaling (*n* = 22), disease (*n* = 55), metabolic (*n* = 14) and regulatory (*n* = 33). According the definitions used in the PW ontology, a “signaling” network comprises mainly of extracellular signal transduction components such as growth factors, kinases, etc. A “regulatory” network, on the other hand, comprises intracellular transcriptional components such as genes, transcription factors, etc. The term “biochemical” here refers to networks that comprises a mix of signaling and regulatory components. “Metabolic” networks consist of components involved in the synthesis and conversion of biomolecules such as enzymes and lipids. Finally, “disease” networks consist of components involved in diseases such as cancer, anemia, pathogenic ailments and disorders such as cell cycle malfunction. Classification 2 was suggested by in-house expertise, where the networks are grouped into four categories (Figure 5(b)), namely metazoan (*n* = 85), cancer (*n* = 24), primitive (*n* = 19) and plants (*n* = 9). The “metazonan” category refers to multicellular organisms and “primitive” refers to unicellular organisms. A given model could naturally belong in multiple categories within a classification but is assigned a unique category for the purpose of simplicity; we chose the categories according to the emphasis laid in the abstracts of the corresponding publications. More details are provided in the SI Appendix (Table S1).

## Results and Discussion

### The regulation of biological models tends to be less nonlinear than expected by chance

The MAE of biological models tends to be less than both the constrained and the unconstrained random ensembles across all approximation orders (Fig 4(a)). At the linear order, for example, the mean MAE of biological models is about 0.025, whereas the mean MAE of the constrained ensembles is twice as large at about 0.05 (*p <* 10^−6^) and even larger for the unconstrained ensembles at about 0.07 (*p <* 10^−23^). This ordering of MAE extends to all approximation orders (Fig 4(a)). Even though the mean MAE of even the unconstrained ensembles tends to be remarkably low (0.07 at worst), the PMAE convey a more accurate sense of approximability. For example, the MAE of the biological models and the constrained ensembles are respectively about 50% and 25% lower than the baseline expectation for the linear order (Fig 4(b)). This means that the biological models are about 25% more linearly approximable than expected by chance, an effect that amplifies with higher approximation orders going up to 50% for order 4, since the PMAE of biological models shrinks faster than the constrained ensembles (Fig 4(b)).

**Figure 4:**
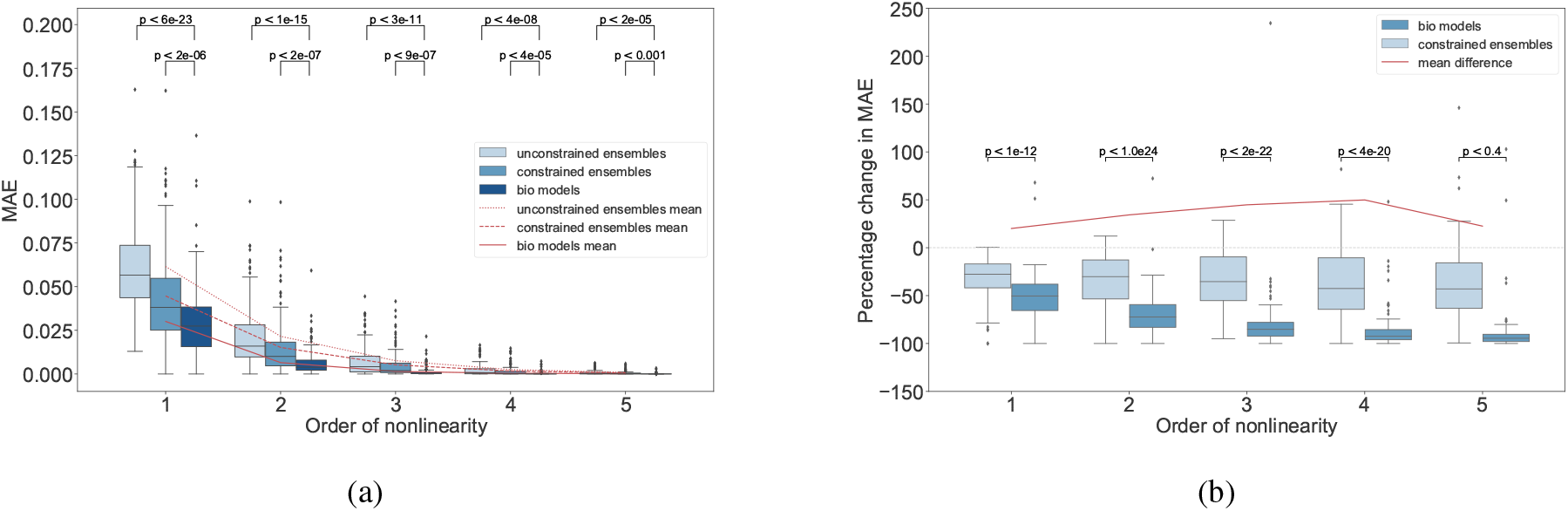
Biological models are more approximable than expected by chance. (a) The MAE of the biological models and the associated constrained and unconstrained ensembles for approximation orders 1 to 5; MAE values for orders 6 and above are negligible and not shown. (b) Percentage change in MAE for the biological models and the associated constrained ensembles computed with respect to the MAE of the corresponding unconstrained ensembles. Every point in the distributions corresponding to the random ensembles represents the average MAE of an ensemble of 100 random networks associated with each biological model. The p-values indicate the statistical significance of the difference in mean MAEs between sets of random ensembles and the biological models for a given order of nonlinearity. Statistical analysis by Welch’s unequal variances t-test.

A possible explanation for the observed trends is that the Taylor spectrums of the biological models tend to be lopsided and more clustered at the linear ends (see Fig 6 for an example), dramatically reducing the load on the higher orders and thus resulting in a faster growth of approximability with higher orders. This effect may be less pronounced in constrained ensembles, and even less in unconstrained ensembles, as the apportioning in the corresponding Taylor spectrums may be less lopsided and more uniform in comparison. Based on these observations we hypothesize, but do not conclude, that biological systems may have been subjected to a slight but discernible selection pressure for developing less nonlinear regulatory rules. This hypothesis is further supported by the fact that the approximability systematically increases as more biological constraints are imposed as one goes from unconstrained to constrained to the biological models (Fig 4). This has implications not only for the feasibility of biomedical approaches to control emergent somatic complexity or guided self-assembly of novel forms [20], but also for models of anatomical homeostasis and evolvability: linearity implies easier control of its own complex processes by any biological system, and more efficient credit assignment during evolution.

### Random models with minimal biological constraints tend to be naturally approximable

The MAE of the unconstrained random ensembles are remarkably low with values averaging around 0.07 and not exceeding 0.17 at the linear order; these values only decrease further with higher approximation orders (Fig 4(a)). These observations suggest that even random Boolean models with minimal biological constraints tend to be considerably approximable. In other words, approximability may be a natural property of random Boolean ensembles and not necessarily a special property of biological models. Furthermore, the MAE has its inflection point at order 2 (Fig 4(a)), meaning that the approximability starts saturating at that order. We hypothesize based on these observations that regulatory information beyond the bilinear order may be inconsequential with regards to the statistics of network dynamics in general.

The above hypothesis has an analogue in the realm of Boolean Ising models, where it was found that maximum entropy (MaxEnt) models with only pairwise interactions were sufficient to fit random multivariate Ising networks that are densely connected and whose state spaces satisfy certain entropy constraints, features that many biological systems share [21]. There are however important differences between our results and theirs: a) even though the Taylor expansions resemble MaxEnt expressions, the latter fit global state-space distributions, whereas our Taylor polynomials are local formulations; and b) whereas our Taylor polynomials are built on derivatives of the states, MaxEnt models are based on the raw values. Notwithstanding these potentially superficial differences, the analogous results are striking and calls for further research on whether the respective explanations are similar if not fundamentally the same. Besides, the above observations are reminiscent of the concept of model degeneracy or ‘sloppiness’, where several models (defined by unique sets of parameters) explain the same biological phenomenon due to redundant parameters [22]. In our case, it’s not the parameters but the higher-order relationships among them that are redundant, as they contribute minimally to the MAE beyond a certain order of approximation (Fig 4(a)). Future research will determine whether there’s a deep connection between the sloppiness of the parameters and the orders of relationships among them.

### The approximability of a biological model depends on its class, with the cancer family displaying the most variability

Even though the nonlinearity of biological networks is less than expected on average, individual and category-dependent variations were observed. In the following, we focus on the “linear approximability” (PMAE corresponding to the linear order) of the biological models since we didn’t observe any order-dependent differences among the models (Figure 4(b)); details are listed in SI Table 1. First, there are a few networks in almost every category that are more nonlinear than expected as evidenced by the negative linear approximability values (Fig 5). Second, the disease networks in C1 and the cancer networks in C2 are the ones with the most linear approximability (high positive values 75%). In other words, the cancer or disease pathways tend to be more optimized for linearity compared to the other categories. This makes sense since a more linear pathway is more amenable to control, which presumably works in favour of the agents of disease. Moreover, the disease and cancer networks also display the highest variability in their linear approximability compared to other categories (*p <* 0.05 in all comparisons for the cancer models), with the PMAE values ranging between -92% and 80%. In other words, disease and cancer models could be either relatively extremely more linear or nonlinear than expected, depending on the specific cases. Note that about 56% of the disease category comprises of non-cancer networks (31/55), which suggests that the effect is not significantly biased by cancer networks. These observations suggest that regulatory nonlinearity may offer an effective “entry point” to the agents of disease by virtue of its natural heterogeneity that they could leverage to their advantage perhaps as a means to evade treatment since there’s no single level of nonlinearity to target. This heterogeneity may also have a connection to one of the hallmarks of cancer, namely genetic heterogeneity [23] where the cancerous cells within an individual display heterogeneous gene expression compared to the homogeneous expression in the healthy cells. In the case of nonlinearity, the heterogeneity manifests at the population level, raising the question of whether it may also be observed at the level of single cells within an individual. In other words, could the heterogeneity of nonlinearity be yet another hallmark of cancer?

**Figure 5:**
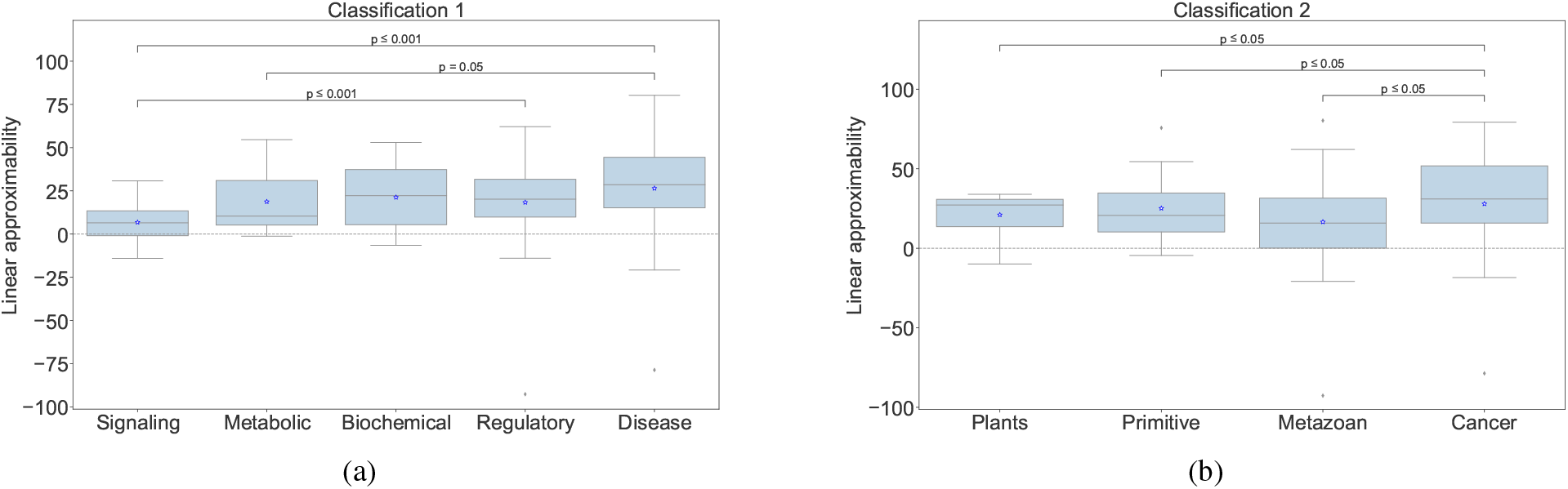
The linear approximabilities of various categories of biological models. (a) Classification 1 (C1); (b) Classification 2 (C2). The categories in either classification are displayed in increasing order of variance. Each box represents the distribution of the linear approximabilities of the corresponding category. The p-values indicate the statistical significance of the difference in the variance between pairs of categories; only the p-values of significantly different pairs are shown. Statistical analysis by F-test of equality of variances.

### The shape of the Taylor spectrum explains the extreme opposite characters of linear approximability of a pair of cancer models

Why are some models more linearly approximable and others less? The answer lies in the organization of the corresponding Taylor decompositions, as described above. To illustrate this in detail, we compared the Taylor decompositions of a pair of models chosen from among the most extreme outliers of linear approximability in either direction (Fig 5). Those models are respectively the following: a linear model describing the role of the protein p53 in the regulation of cell-cycle arrest in breast cancer [24] and henceforth referred to as the ‘P53’ model; and a nonlinear model describing the role of mutations in the regulation of metastasis in lung cancer [25] and henceforth referred to as the ‘Metastasis’ model. P53 has a linear approximability of about 72%, and it consists of 16 nodes with a mean in-degree of 3.8 ± 2.4 and a mean output bias of 0.38 ± 0.14; and, Metastasis has a linear approximability of about -79%, and it consists of 32 nodes with a mean in-degree of 4.9 ± 2.5 and a mean output bias of 0.27 ± 0.26. Thus, while P53 is smaller and sparser than Metastasis, its nodes exhibit more output-uncertainty compared to Metastasis. According to the mean field theory of random Boolean networks [26], the opposite characters of the mean in-degree and the output bias of these models means that their dynamical behaviors could be expected to be similar (although with the caution that the theory was originally developed for infinite-sized and homogeneously connected networks, which is not the case here). However, we know that their linear approximabilities, which is another expression of dynamical behavior, are opposites. One explanation for this discrepancy lies in the distinct apportioning of nonlinearity in their respective

Taylor decompositions (Fig 6). Specifically, while the magnitude of nonlinearity, defined as the mean absolute value of the Taylor derivatives for a given order and normalized appropriately (see text of Fig 6), tends to be clustered around the linear order for P53, they are relatively more spread out for Metastasis. Moreover, while the magnitude of the linear order for P53 is more than twice as large the next largest magnitude at lower orders the corresponding ratios for Metastasis are relatively smaller, thus explaining why P53 is more linearly approximable than Metastasis. This result is consistent with predictions based on a model of scaling of cellular control policies [27]. A more controllable (linear) network (P53) is optimal for cooperation with other cells towards collective (normal morphogenetic) goals. In contrast, a cell defecting from the collective and reverting to a more unicellular lifestyle (Metastasis) should exhibit a less predictable, controllable network due to pressures from parasites and competitors that independent unicellular organisms face. Methods for calculating controllability (e.g., linearity) are an important addition to recent efforts to solve the conundrum of interpretability of information structures in contexts ranging from machine learning to evolutionary developmental biology [28, 29, 30].

**Figure 6:**
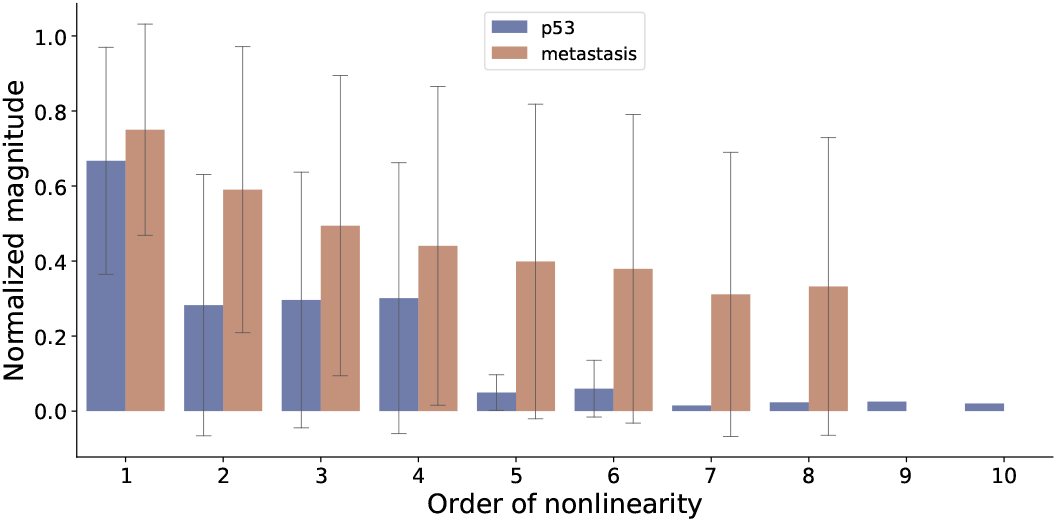
A comparison of the spectrums of the magnitudes of nonlinearity of the models corresponding to the two extreme cases of approximability. The bars represent the mean absolute value of all the Taylor derivatives of the corresponding order, averaged over all the nodes of the corresponding model containing those terms. The maximum orders (in-degrees) of P53 and Metastasis are respectively 10 and 8. and To compare the derivatives from different models they were normalized with respect to the maximum possible absolute value of a Taylor derivative of order |*α*| (Equation 1) of a Boolean function with output bias *p*, given by (min(*p*, 0.5) − max(*p* − 0.5, 0))2^|*α*|^ (proof provided in the SI Appendix). The error bars represent the standard deviation and not the confidence intervals of the means since they are not estimates. The errors appear large whenever the corresponding distribution of the magnitudes are bimodal with many values clustered close to 0.

### Broader implications

This paper introduces the concept of regulatory nonlinearity as a characteristic of Boolean networks. There are several other related characterizations of Boolean networks such as canalization [31], effective connectivity [11], symmetry [32] and controllability [33]. It has been previously reported that the levels of canalization (a measure of the extent to which fewer inputs influence the outputs of a Boolean function) and the mean effective connectivity (a measure of collective canalization) are high in biological networks [3, 11]. It has also been found that biological networks need few inputs to reprogram [34] and are relatively easier to control [4]. Our formulation of regulatory nonlinearity is related to these other measures in that more linearity implies more apportioning of influence to individual inputs rather than collective sets of inputs (Figure 6). Hence, we hypothesize that regulatory nonlinearity may serve the purpose of controllability and epigenetic stability [35].

Our results further moot the possibility that regulatory nonlinearity may be a factor underlying more powerful dynamical phenomena such as memory [36] and computation, defined as the capacity for adaptive information-processing [37]. Even though there’s increasing consensus that biological systems contain memory and perform computation, clarity is lacking as to what features of those systems enable it and what general principles underlie it [37, 38]. Our framework of regulatory nonlinearity offers an approach to answering these questions. For example, one could consider a known dynamical model with a capacity for memory [36] or universal computation such as the elementary cellular automaton (ECA) driven by rule 110 [39] and ask if there are unique properties of its Taylor spectrum that confer their respective capabilities. Present approaches to answering this question typically consists of characterization of the dynamical behavior and not the rules [40, 41, 36]. A characterization of the rules especially makes sense for ECA [42] since the structure is always the same (lattice) and the only feature that distinguishes one ECA from the other is the rule.

Looking at such questions from an even broader perspective it becomes evident that they are only instances of the ultimate puzzle of complex systems, namely what connects the structure and the function of a system. Even though recent work has attempted to answer this question from the perspective of the rules or dynamical laws that govern the system [43, 11, 42], more tools are needed [44]. In that regard, our framework of regulatory nonlinearity could be a novel addition to this burgeoning toolkit in that it could also be applied to continuous differentiable models of biological networks such as those based on differential equations.

### Limitations

The main limitation of our formulation of approximability is that the approximation accuracy will necessarily increase with higher orders of approximation since the Taylor terms would be accumulated with each higher order (the highest order of approximation is exact). However, this does not affect the falsifiability of our framework since it’s possible, for example, to construct networks with XOR-like functions that would be clearly less linearly approximable than the associated ensembles. The Metastasis model is another example in that regard (Figure 6). Furthermore, though our experiments control for regulatory nonlinearity they don’t offer insights into how it dynamically interacts with other network features such as network structure in generating the observed approximability. For example, the network structure may be expected to determine which states the network enters into at any given point that in turn determines which parts of the Taylor compositions are actually utilized and how. Moreover, it’s known that certain characteristics of the local regulatory rules, such as “effective connectivity”, dynamically modulate the network structure itself [45]. Thus, more research is required to investigate the extent to which the structure and output bias facilitates the high approximability of biological networks and vice versa, for which we already have produced preliminary results by way of the constrained models (Fig 4). Lastly, our conclusions about the linearity of biological regulatory networks may be a reflection of a hidden bias built in the inference methods that produced the models in the first place. We leave it to future work to explore these realms.

## 2 Data Availability

We used 137 Boolean network models of gene regulation from two sources, namely the *cell collective* [9] and reference [3]. Subsequently, we categorized these models using two types of classifications C1 and C2 as described in Section 1. We provide the details of the classification of each model along with their Pudmed ID and their linear approximation score in the supplementary information file (SI), Table S1.

## 3 Code Availability

The code that we used for the approximations and the simulations are available through this link: https://gitlab.com/smanicka/boolion.

## 4 Author Contributions

S.M. and D.M. conceived the study; S.M. and K.J. designed code; K.J. performed research and simulations; S.M., K.J., and D.M. analyzed data; M.L. helped bridge the theory to biology. All authors helped in the writing of the manuscript. All authors approved the final version of the manuscript.

## 5 Acknowledgements

We thank Claus Kadelka for sharing a large repository of Boolean models curated by his group and the accompanying *Python* code for analyzing it. D.M. was partially supported by a Collaboration grant (850896) from the Simons Foundation. M.L. gratefully acknowledges support via grant TWCF0606 of the Templeton World Charity Foundation.

## Supplementary Information

### 1. Probabilistic generalization of Boolean logic

Here we show that 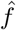 is a generalization of *f* in the sense that 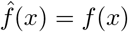.*x*∈{0,1}^n^

#### Theorem 0.1.

*For discrete values of x*_*i*_ ∈ {0, 1}, *i* = 1, …, *n, we have*

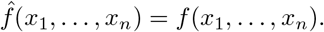

*Proof*. Let *z* = (*z*_1_, …, *z*_*n*_) ∈ {0, 1}^*n*^. Since each *z*_*i*_ is either 0 or 1, we have that *p*_*i*_ = 1 if *z*_*i*_ = 1 or *p*_*i*_ = 0 if *z*_*i*_ = 0 for *i* = 1, …, *n*. We want to show that 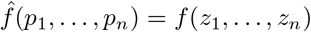. Since 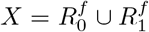, we have that either 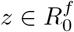 or 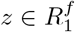. If 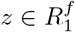, then *f* (*z*) = 1 and 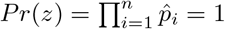. Moreover, for any other 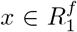 with *x* ≠ *z* we have that *Pr*(*x*) = 0. Thus, 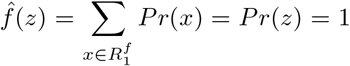. Now if 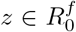, then 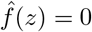 because Σ_∅_ = 0. Thus, 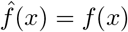 for all *x* ∈ {0, 1}^*n*^.

### 2. Maximum absolute value of a Taylor derivative

Here we show that max 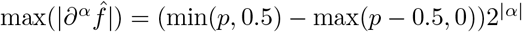. We begin with the definition of the derivative given by

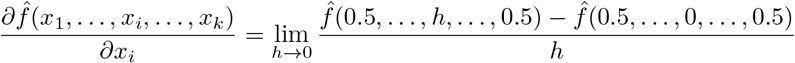

Since 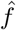 is a pseudo-Boolean function and hence a multilinear polynomial [1], we can rewrite it by setting *h* to 1, as a finite difference; the idea being that the derivative taken over any point on a line is the line itself:

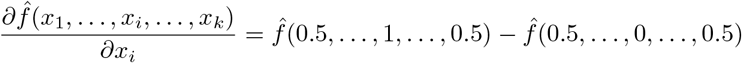

Since multilinear interpolation can be formulated as weighted averaging [2], we can further rewrite it as follows (the weights are equal here since the non-binary values are all set to 0.5):

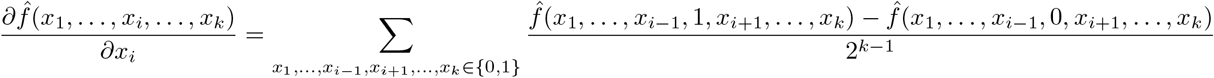

As can be seen, there are a total of 2^*k*^ terms, with half of them positively signed and half negative. This form of expression generalizes to derivatives taken over two or more variables. For example, the derivative taken over two variables, *x*_*i*_ and *x*_*j*_, looks as follows:

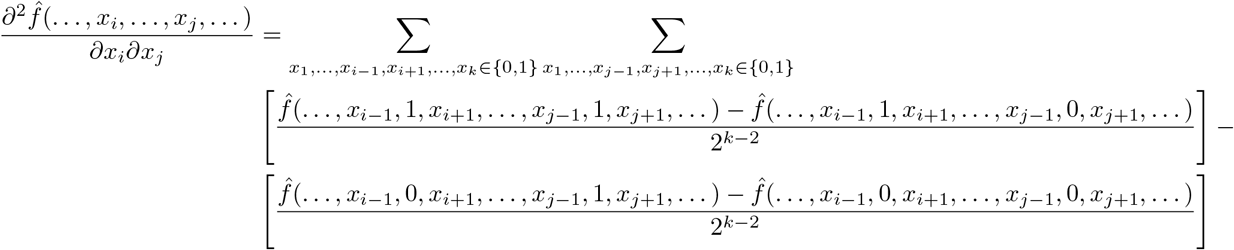

Following rearrangement of terms it becomes evident that this expression also contains 2^*k*−1^ positive terms and 2^*k*−1^ negative terms, with the only difference in the power of the denominator term. It can thus be concluded that any derivative 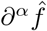 (in multi-index notation) has 2^*k*−1^ positive terms and 2^*k*−1^ negative terms. A straightforward way to maximize the value of a derivative expressed in this form is by assigning as many instances of 1 as possible to the positive terms and as few instances of 1 as possible to the negative terms. For a Boolean function with *k* inputs and output bias *p*, this can be accomplished by assigning min(2^*k*−1^, *p*2^*k*^) ones and max(*p*2^*k*^ − 2^*k*−1^, 0) ones respectively. Therefore,

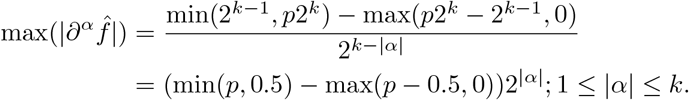

Note that this formula only applies to a specific order of nonlinearity |*α*| independent of the other orders within the same Boolean function. In actuality, there are dependencies between the various orders within a Boolean function. That is, if a Boolean function were to be constructed such that the derivative of a particular order |*α*_1_| is maximized then there’s no guarantee that the derivative of another order |*α*_2_| = |*α*_1_| could be simultaneously maximized. This is one of the limitations of the normalization for which the above formula is used.

**Table 1:**
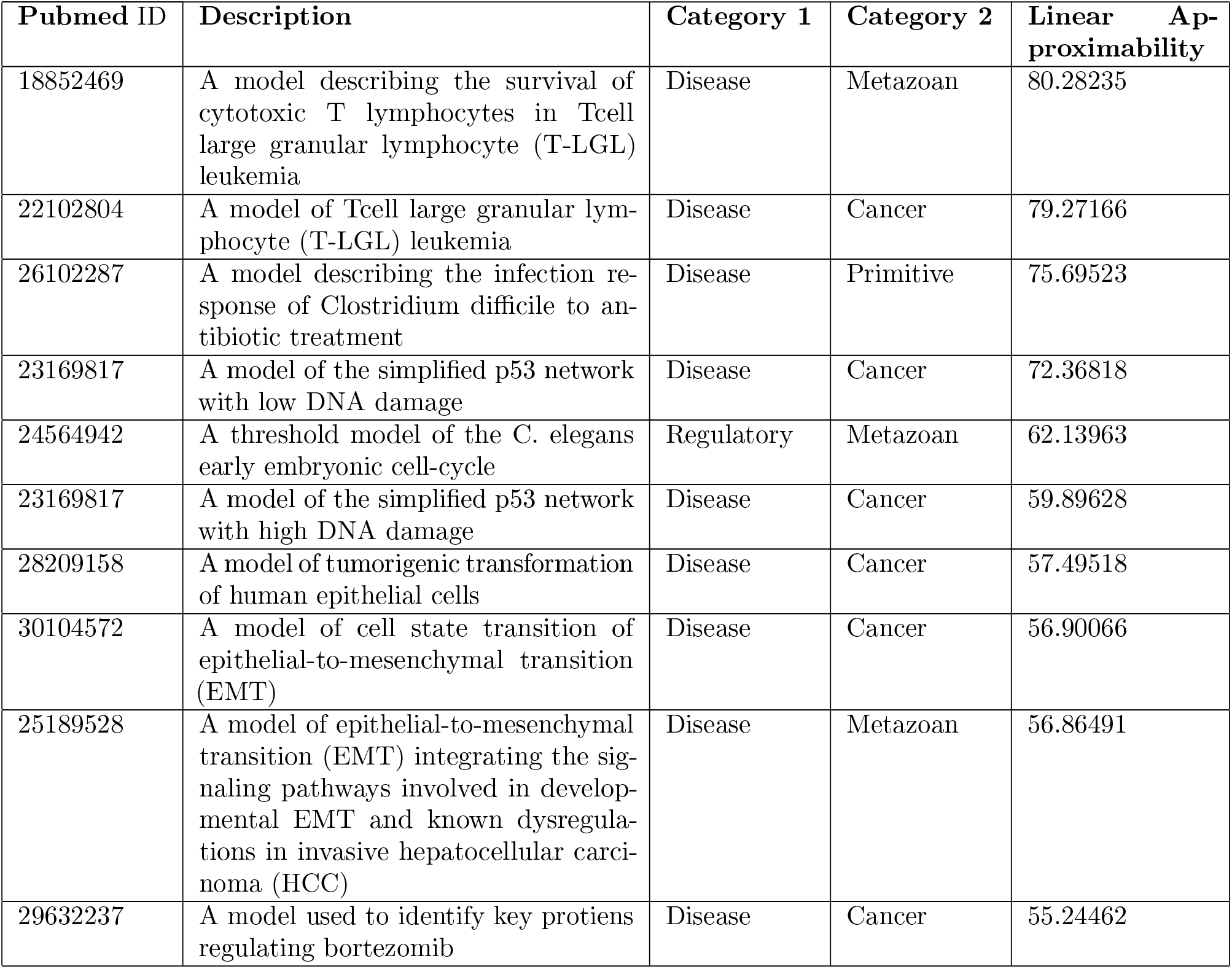
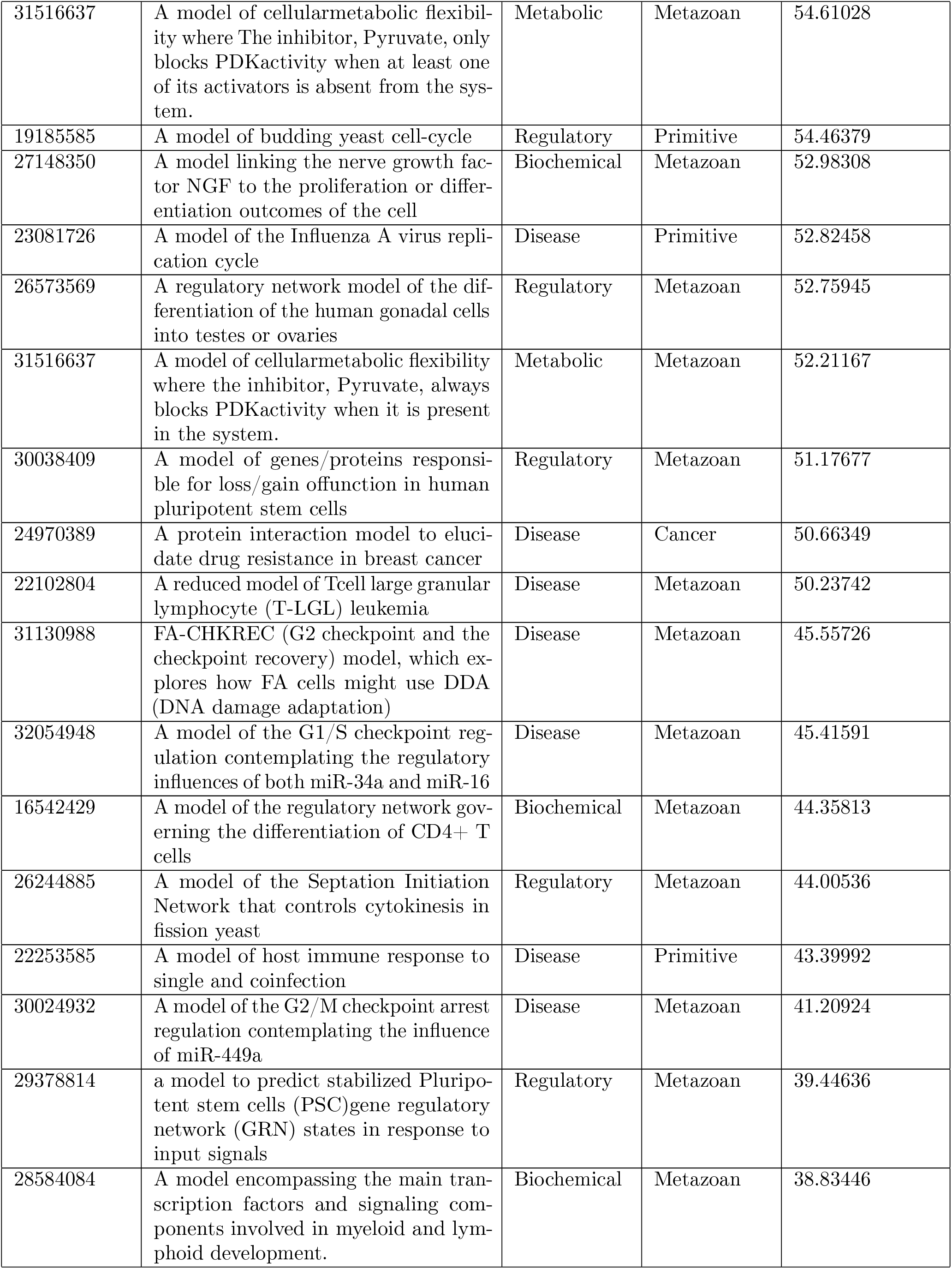
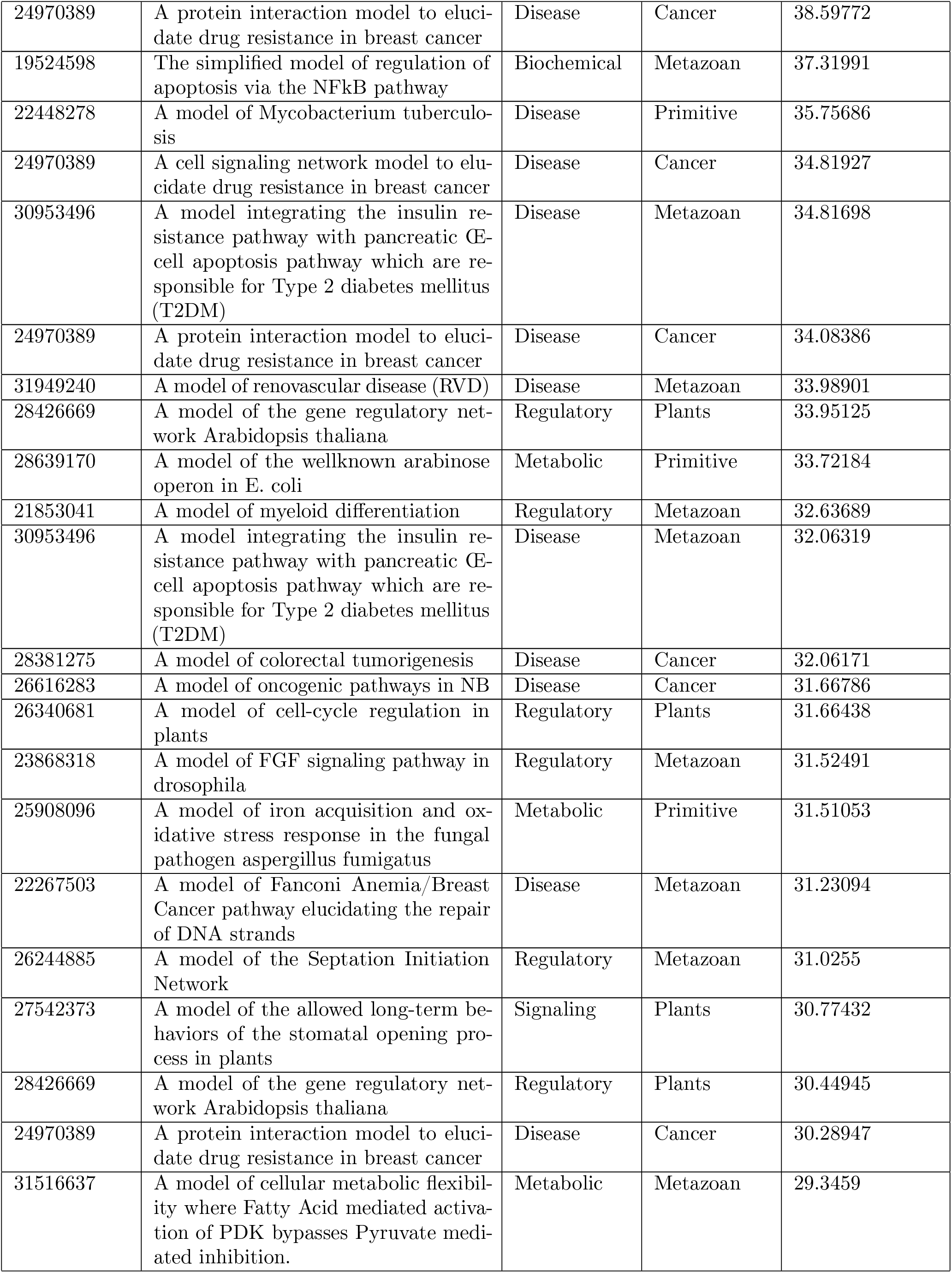
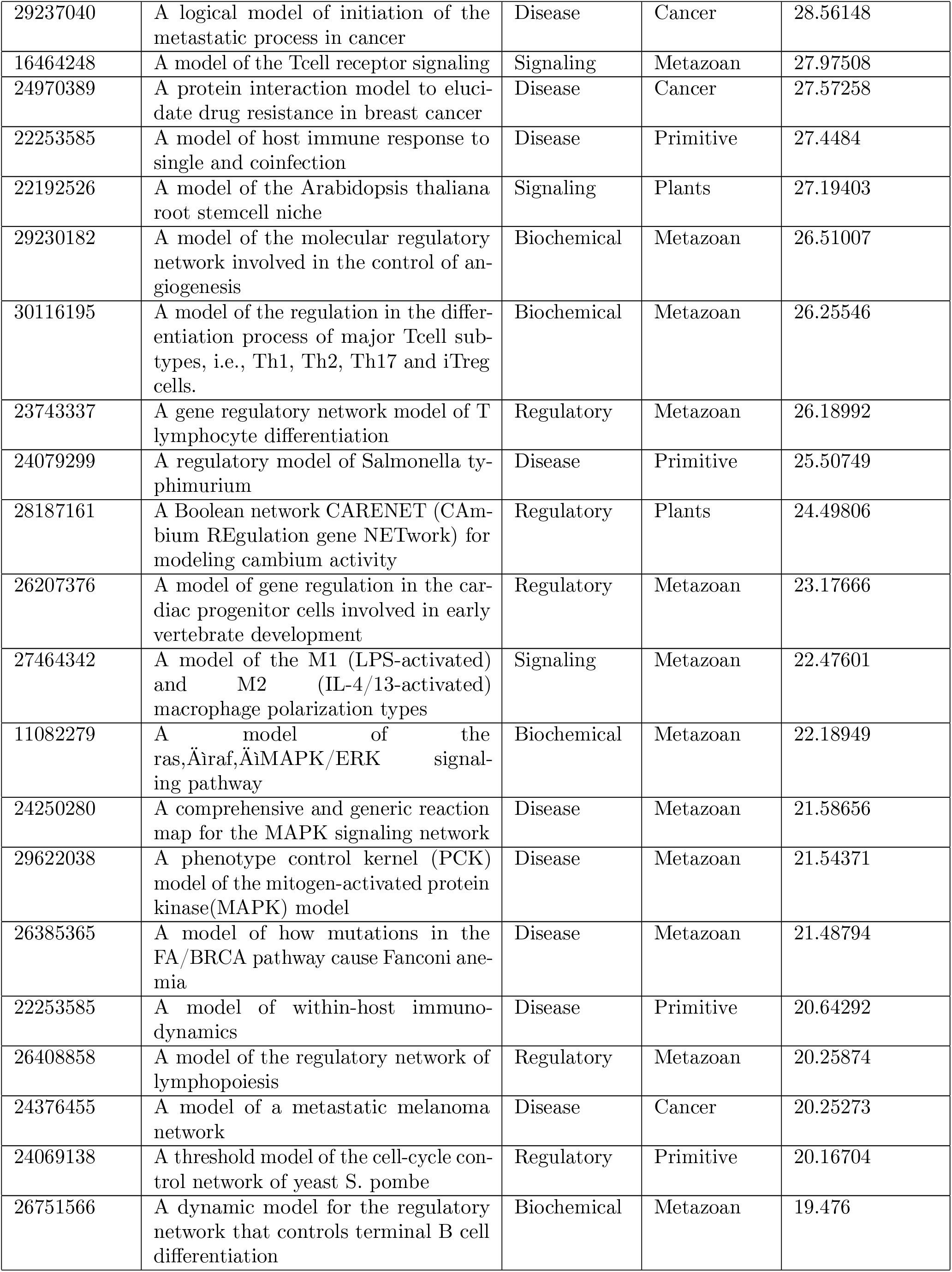
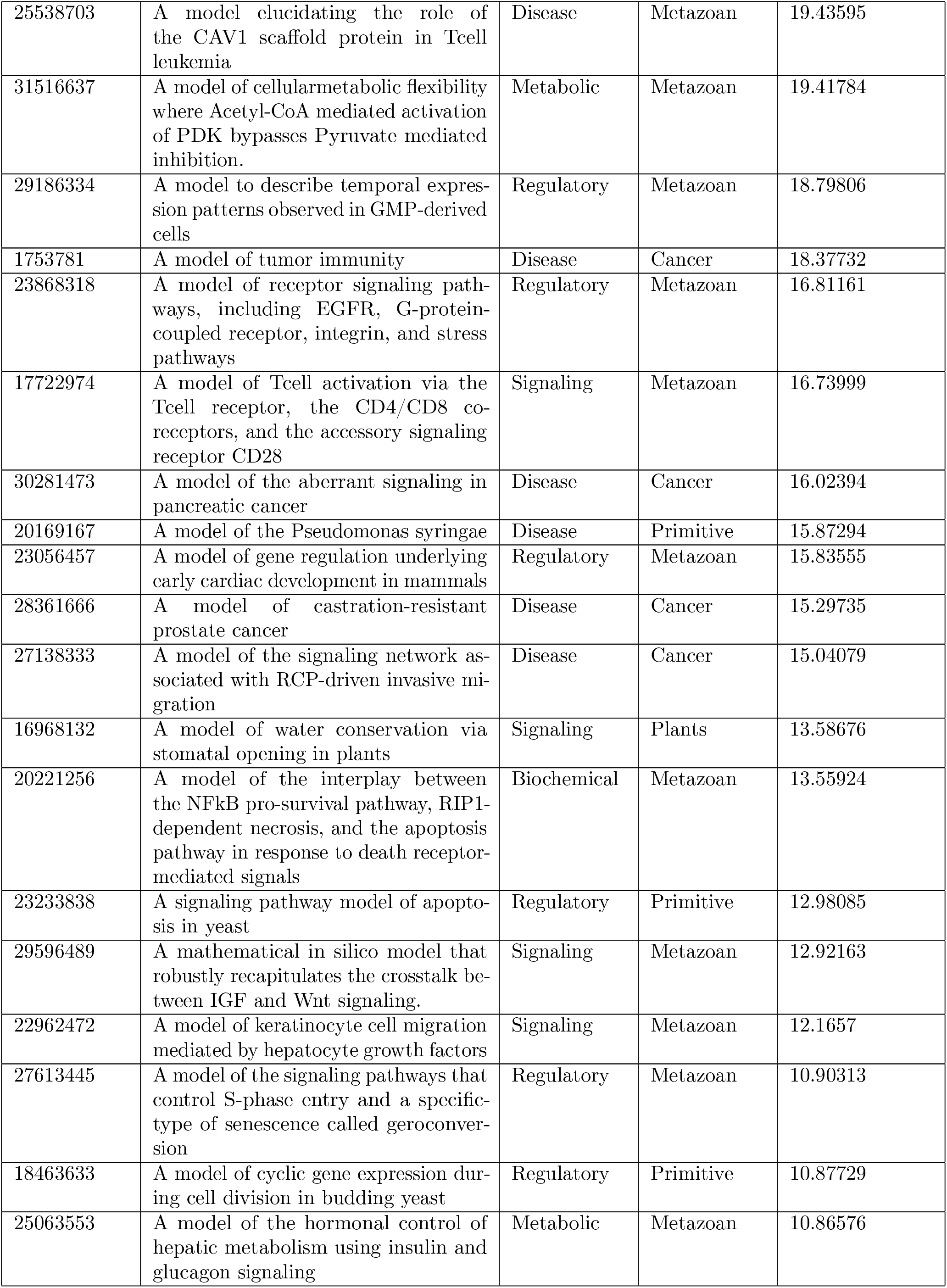
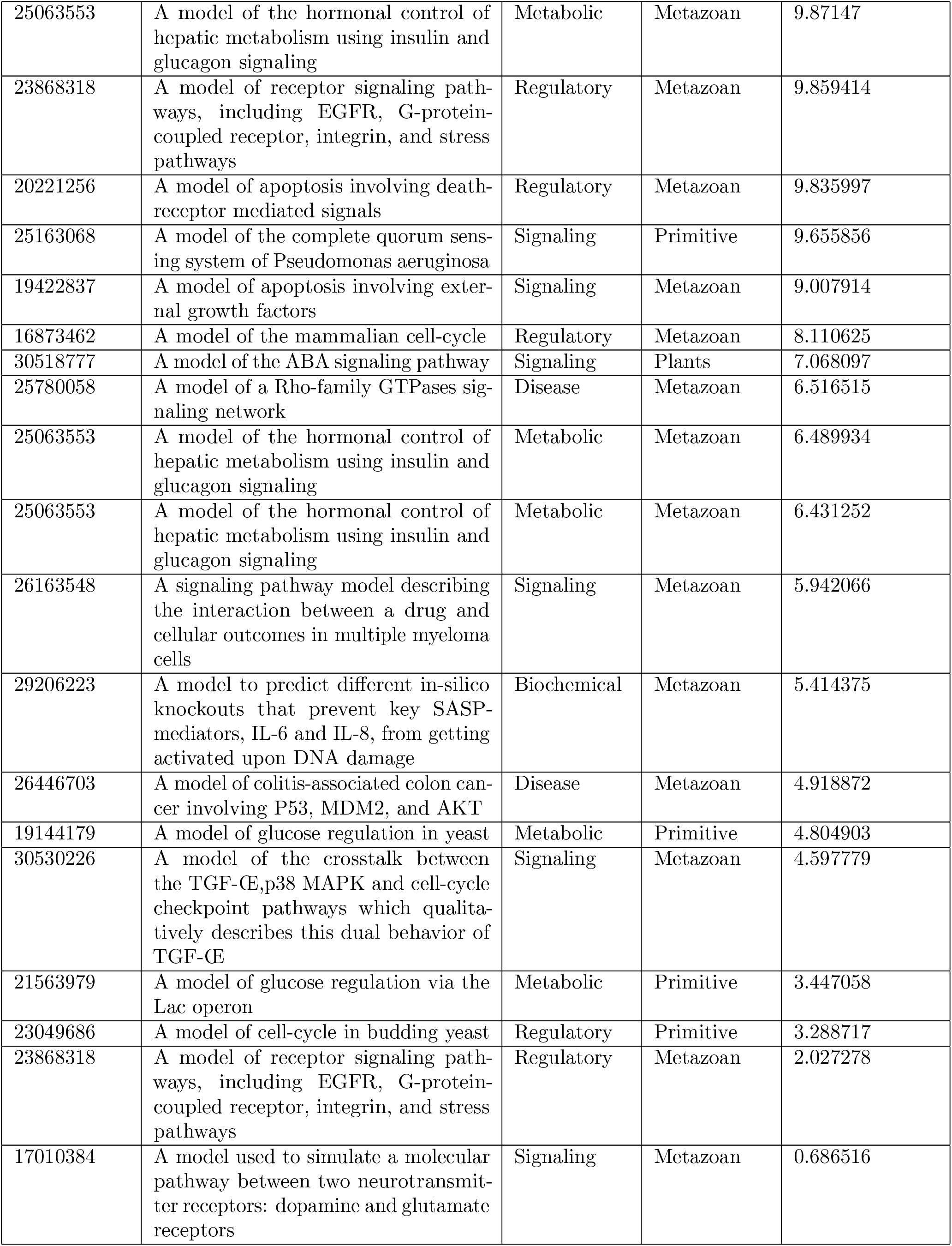
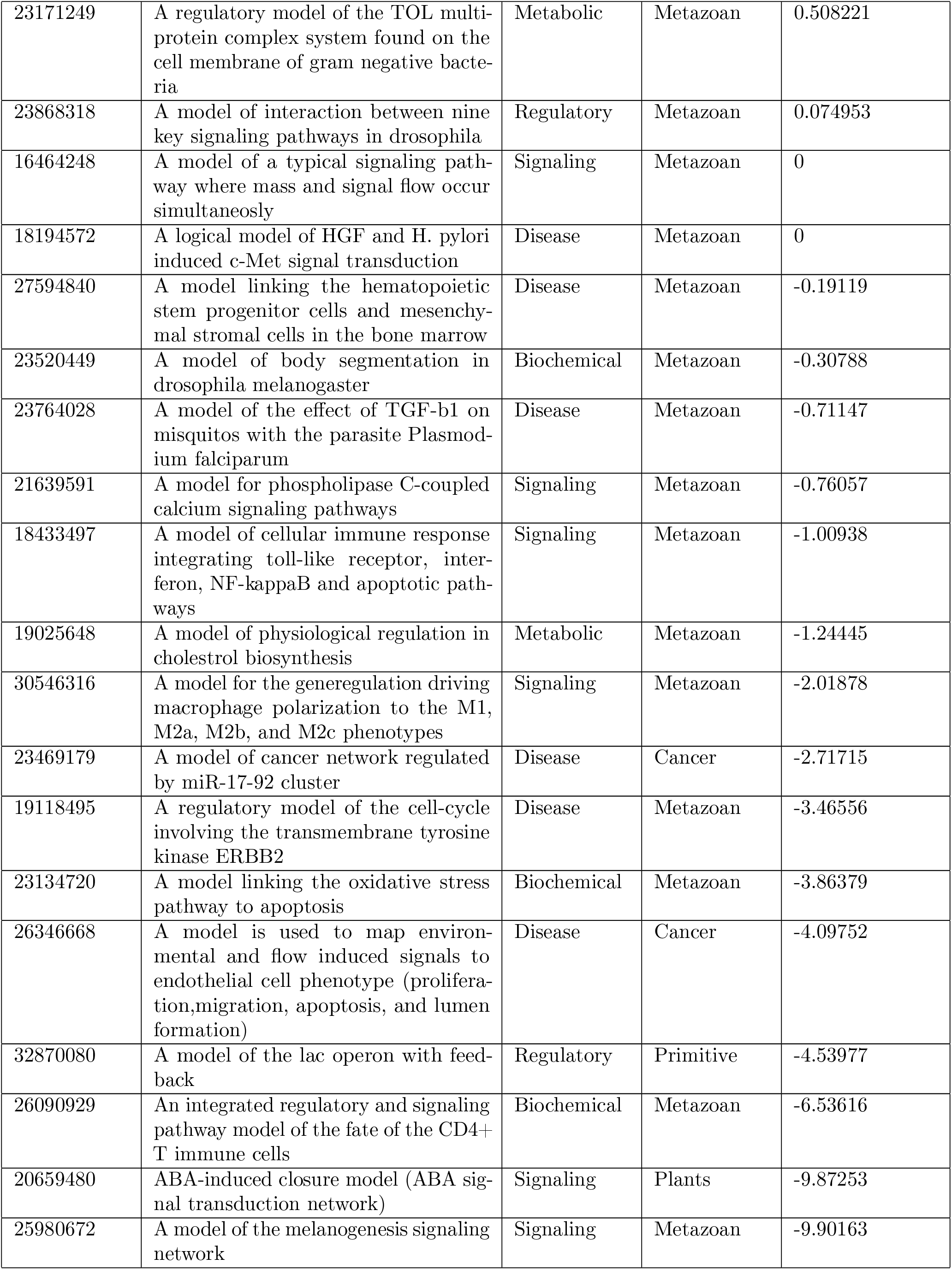
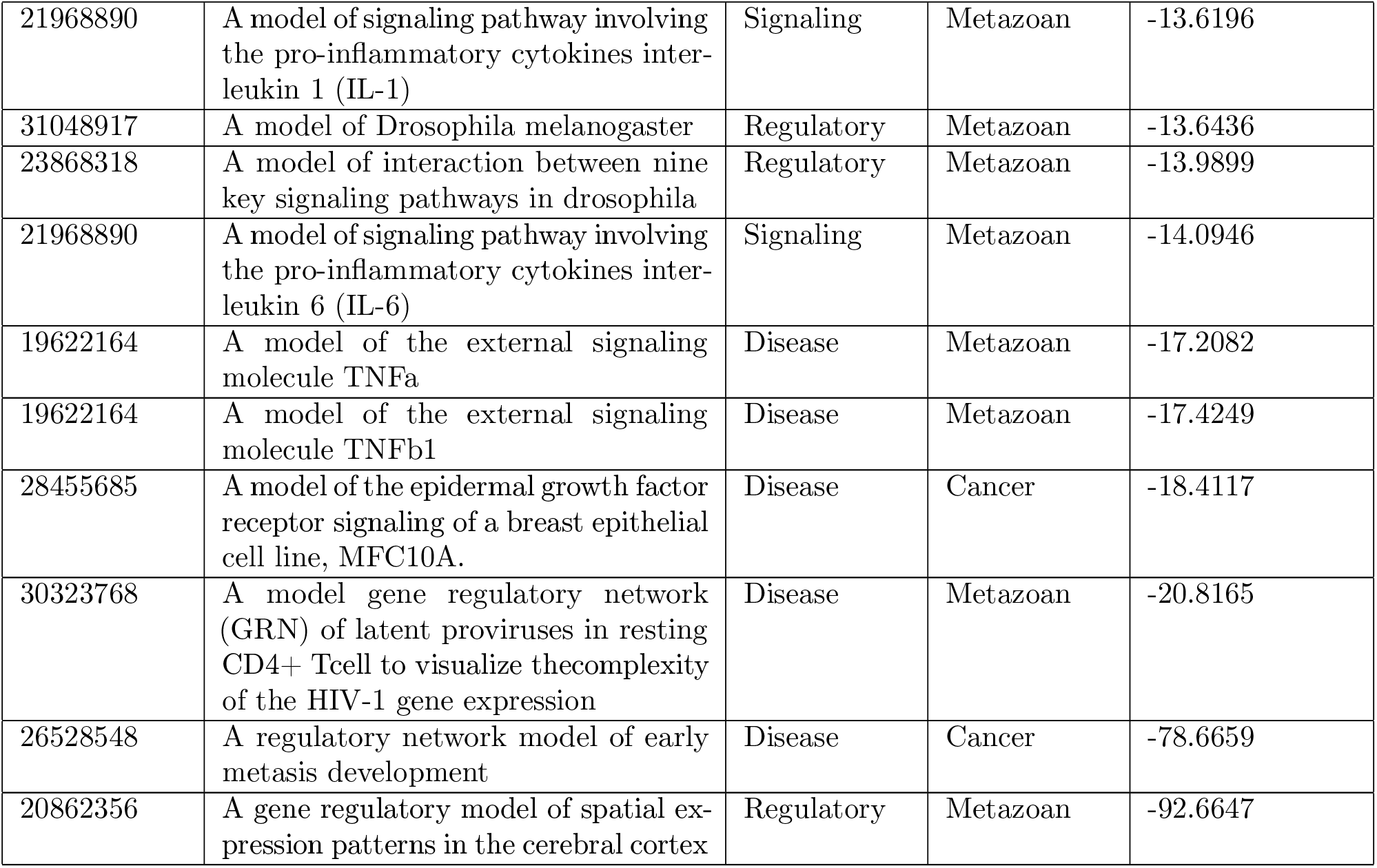
Dataset Description

| Pubmed ID | Description | Category 1 | Category 2 | Linear Ap-<br>proximability |
| --- | --- | --- | --- | --- |
| 18852469 | A model describing the survival of cytotoxic T lymphocytes in Tcell large granular lymphocyte (T-LGL) leukemia | Disease | Metazoan | 80.28235 |
| 22102804 | A model of Tcell large granular lymphocyte (T-LGL) leukemia | Disease | Cancer | 79.27166 |
| 26102287 | A model describing the infection response of Clostridium difficile to antibiotic treatment | Disease | Primitive | 75.69523 |
| 23169817 | A model of the simplified p53 network with low DNA damage | Disease | Cancer | 72.36818 |
| 24564942 | A threshold model of the C. elegans early embryonic cell-cycle | Regulatory | Metazoan | 62.13963 |
| 23169817 | A model of the simplified p53 network with high DNA damage | Disease | Cancer | 59.89628 |
| 28209158 | A model of tumorigenic transformation of human epithelial cells | Disease | Cancer | 57.49518 |
| 30104572 | A model of cell state transition of epithelial-to-mesenchymal transition (EMT) | Disease | Cancer | 56.90066 |
| 25189528 | A model of epithelial-to-mesenchymal transition (EMT) integrating the signaling pathways involved in developmental EMT and known dysregulations in invasive hepatocellular carcinoma (HCC) | Disease | Metazoan | 56.86491 |
| 29632237 | A model used to identify key proteins regulating bortezomib | Disease | Cancer | 55.24462 |
| 31516637 | A model of cellular metabolic flexibility where The inhibitor, Pyruvate, only blocks PDK activity when at least one of its activators is absent from the system. | Metabolic | Metazoan | 54.61028 |
| 19185585 | A model of budding yeast cell-cycle | Regulatory | Primitive | 54.46379 |
| 27148350 | A model linking the nerve growth factor NGF to the proliferation or differentiation outcomes of the cell | Biochemical | Metazoan | 52.98308 |
| 23081726 | A model of the Influenza A virus replication cycle | Disease | Primitive | 52.82458 |
| 26573569 | A regulatory network model of the differentiation of the human gonadal cells into testes or ovaries | Regulatory | Metazoan | 52.75945 |
| 31516637 | A model of cellular metabolic flexibility where the inhibitor, Pyruvate, always blocks PDK activity when it is present in the system. | Metabolic | Metazoan | 52.21167 |
| 30038409 | A model of genes/proteins responsible for loss/gain of function in human pluripotent stem cells | Regulatory | Metazoan | 51.17677 |
| 24970389 | A protein interaction model to elucidate drug resistance in breast cancer | Disease | Cancer | 50.66349 |
| 22102804 | A reduced model of T cell large granular lymphocyte (T-LGL) leukemia | Disease | Metazoan | 50.23742 |
| 31130988 | FA-CHKREC (G2 checkpoint and the checkpoint recovery) model, which explores how FA cells might use DDA (DNA damage adaptation) | Disease | Metazoan | 45.55726 |
| 32054948 | A model of the G1/S checkpoint regulation contemplating the regulatory influences of both miR-34a and miR-16 | Disease | Metazoan | 45.41591 |
| 16542429 | A model of the regulatory network governing the differentiation of CD4+ T cells | Biochemical | Metazoan | 44.35813 |
| 26244885 | A model of the Septation Initiation Network that controls cytokinesis in fission yeast | Regulatory | Metazoan | 44.00536 |
| 22253585 | A model of host immune response to single and coinfection | Disease | Primitive | 43.39992 |
| 30024932 | A model of the G2/M checkpoint arrest regulation contemplating the influence of miR-449a | Disease | Metazoan | 41.20924 |
| 29378814 | a model to predict stabilized Pluripotent stem cells (PSC) gene regulatory network (GRN) states in response to input signals | Regulatory | Metazoan | 39.44636 |
| 28584084 | A model encompassing the main transcription factors and signaling components involved in myeloid and lymphoid development. | Biochemical | Metazoan | 38.83446 |
| 24970389 | A protein interaction model to elucidate drug resistance in breast cancer | Disease | Cancer | 38.59772 |
| 19524598 | The simplified model of regulation of apoptosis via the NFkB pathway | Biochemical | Metazoan | 37.31991 |
| 22448278 | A model of Mycobacterium tuberculosis | Disease | Primitive | 35.75686 |
| 24970389 | A cell signaling network model to elucidate drug resistance in breast cancer | Disease | Cancer | 34.81927 |
| 30953496 | A model integrating the insulin resistance pathway with pancreatic E-cell apoptosis pathway which are responsible for Type 2 diabetes mellitus (T2DM) | Disease | Metazoan | 34.81698 |
| 24970389 | A protein interaction model to elucidate drug resistance in breast cancer | Disease | Cancer | 34.08386 |
| 31949240 | A model of renovascular disease (RVD) | Disease | Metazoan | 33.98901 |
| 28426669 | A model of the gene regulatory network Arabidopsis thaliana | Regulatory | Plants | 33.95125 |
| 28639170 | A model of the wellknown arabinose operon in E. coli | Metabolic | Primitive | 33.72184 |
| 21853041 | A model of myeloid differentiation | Regulatory | Metazoan | 32.63689 |
| 30953496 | A model integrating the insulin resistance pathway with pancreatic E-cell apoptosis pathway which are responsible for Type 2 diabetes mellitus (T2DM) | Disease | Metazoan | 32.06319 |
| 28381275 | A model of colorectal tumorigenesis | Disease | Cancer | 32.06171 |
| 26616283 | A model of oncogenic pathways in NB | Disease | Cancer | 31.66786 |
| 26340681 | A model of cell-cycle regulation in plants | Regulatory | Plants | 31.66438 |
| 23868318 | A model of FGF signaling pathway in drosophila | Regulatory | Metazoan | 31.52491 |
| 25908096 | A model of iron acquisition and oxidative stress response in the fungal pathogen aspergillus fumigatus | Metabolic | Primitive | 31.51053 |
| 22267503 | A model of Fanconi Anemia/Breast Cancer pathway elucidating the repair of DNA strands | Disease | Metazoan | 31.23094 |
| 26244885 | A model of the Septation Initiation Network | Regulatory | Metazoan | 31.0255 |
| 27542373 | A model of the allowed long-term behaviors of the stomatal opening process in plants | Signaling | Plants | 30.77432 |
| 28426669 | A model of the gene regulatory network Arabidopsis thaliana | Regulatory | Plants | 30.44945 |
| 24970389 | A protein interaction model to elucidate drug resistance in breast cancer | Disease | Cancer | 30.28947 |
| 31516637 | A model of cellular metabolic flexibility where Fatty Acid mediated activation of PDK bypasses Pyruvate mediated inhibition. | Metabolic | Metazoan | 29.3459 |
| 29237040 | A logical model of initiation of the metastatic process in cancer | Disease | Cancer | 28.56148 |
| 16464248 | A model of the Tcell receptor signaling | Signaling | Metazoan | 27.97508 |
| 24970389 | A protein interaction model to elucidate drug resistance in breast cancer | Disease | Cancer | 27.57258 |
| 22253585 | A model of host immune response to single and coinfection | Disease | Primitive | 27.4484 |
| 22192526 | A model of the Arabidopsis thaliana root stemcell niche | Signaling | Plants | 27.19403 |
| 29230182 | A model of the molecular regulatory network involved in the control of angiogenesis | Biochemical | Metazoan | 26.51007 |
| 30116195 | A model of the regulation in the differentiation process of major Tcell subtypes, i.e., Th1, Th2, Th17 and iTreg cells. | Biochemical | Metazoan | 26.25546 |
| 23743337 | A gene regulatory network model of T lymphocyte differentiation | Regulatory | Metazoan | 26.18992 |
| 24079299 | A regulatory model of Salmonella typhimurium | Disease | Primitive | 25.50749 |
| 28187161 | A Boolean network CARENET (C**A**m**B**ium **R**Egulation gene **N**ETwork) for modeling cambium activity | Regulatory | Plants | 24.49806 |
| 26207376 | A model of gene regulation in the cardiac progenitor cells involved in early vertebrate development | Regulatory | Metazoan | 23.17666 |
| 27464342 | A model of the M1 (LPS-activated) and M2 (IL-4/13-activated) macrophage polarization types | Signaling | Metazoan | 22.47601 |
| 11082279 | A model of the ras,Äiraf,ÄiMAPK/ERK signaling pathway | Biochemical | Metazoan | 22.18949 |
| 24250280 | A comprehensive and generic reaction map for the MAPK signaling network | Disease | Metazoan | 21.58656 |
| 29622038 | A phenotype control kernel (PCK) model of the mitogen-activated protein kinase(MAPK) model | Disease | Metazoan | 21.54371 |
| 26385365 | A model of how mutations in the FA/BRCA pathway cause Fanconi anemia | Disease | Metazoan | 21.48794 |
| 22253585 | A model of within-host immunodynamics | Disease | Primitive | 20.64292 |
| 26408858 | A model of the regulatory network of lymphopoiesis | Regulatory | Metazoan | 20.25874 |
| 24376455 | A model of a metastatic melanoma network | Disease | Cancer | 20.25273 |
| 24069138 | A threshold model of the cell-cycle control network of yeast *S. pombe* | Regulatory | Primitive | 20.16704 |
| 26751566 | A dynamic model for the regulatory network that controls terminal B cell differentiation | Biochemical | Metazoan | 19.476 |
| 25538703 | A model elucidating the role of the CAV1 scaffold protein in Tcell leukemia | Disease | Metazoan | 19.43595 |
| 31516637 | A model of cellularmetabolic flexibility where Acetyl-CoA mediated activation of PDK bypasses Pyruvate mediated inhibition. | Metabolic | Metazoan | 19.41784 |
| 29186334 | A model to describe temporal expression patterns observed in GMP-derived cells | Regulatory | Metazoan | 18.79806 |
| 1753781 | A model of tumor immunity | Disease | Cancer | 18.37732 |
| 23868318 | A model of receptor signaling pathways, including EGFR, G-protein-coupled receptor, integrin, and stress pathways | Regulatory | Metazoan | 16.81161 |
| 17722974 | A model of Tcell activation via the Tcell receptor, the CD4/CD8 co-receptors, and the accessory signaling receptor CD28 | Signaling | Metazoan | 16.73999 |
| 30281473 | A model of the aberrant signaling in pancreatic cancer | Disease | Cancer | 16.02394 |
| 20169167 | A model of the Pseudomonas syringae | Disease | Primitive | 15.87294 |
| 23056457 | A model of gene regulation underlying early cardiac development in mammals | Regulatory | Metazoan | 15.83555 |
| 28361666 | A model of castration-resistant prostate cancer | Disease | Cancer | 15.29735 |
| 27138333 | A model of the signaling network associated with RCP-driven invasive migration | Disease | Cancer | 15.04079 |
| 16968132 | A model of water conservation via stomatal opening in plants | Signaling | Plants | 13.58676 |
| 20221256 | A model of the interplay between the NFkB pro-survival pathway, RIP1-dependent necrosis, and the apoptosis pathway in response to death receptor-mediated signals | Biochemical | Metazoan | 13.55924 |
| 23233838 | A signaling pathway model of apoptosis in yeast | Regulatory | Primitive | 12.98085 |
| 29596489 | A mathematical in silico model that robustly recapitulates the crosstalk between IGF and Wnt signaling. | Signaling | Metazoan | 12.92163 |
| 22962472 | A model of keratinocyte cell migration mediated by hepatocyte growth factors | Signaling | Metazoan | 12.1657 |
| 27613445 | A model of the signaling pathways that control S-phase entry and a specific-type of senescence called geroconversion | Regulatory | Metazoan | 10.90313 |
| 18463633 | A model of cyclic gene expression during cell division in budding yeast | Regulatory | Primitive | 10.87729 |
| 25063553 | A model of the hormonal control of hepatic metabolism using insulin and glucagon signaling | Metabolic | Metazoan | 10.86576 |
| 25063553 | A model of the hormonal control of hepatic metabolism using insulin and glucagon signaling | Metabolic | Metazoan | 9.87147 |
| 23868318 | A model of receptor signaling pathways, including EGFR, G-protein-coupled receptor, integrin, and stress pathways | Regulatory | Metazoan | 9.859414 |
| 20221256 | A model of apoptosis involving death-receptor mediated signals | Regulatory | Metazoan | 9.835997 |
| 25163068 | A model of the complete quorum sensing system of *Pseudomonas aeruginosa* | Signaling | Primitive | 9.655856 |
| 19422837 | A model of apoptosis involving external growth factors | Signaling | Metazoan | 9.007914 |
| 16873462 | A model of the mammalian cell-cycle | Regulatory | Metazoan | 8.110625 |
| 30518777 | A model of the ABA signaling pathway | Signaling | Plants | 7.068097 |
| 25780058 | A model of a Rho-family GTPases signaling network | Disease | Metazoan | 6.516515 |
| 25063553 | A model of the hormonal control of hepatic metabolism using insulin and glucagon signaling | Metabolic | Metazoan | 6.489934 |
| 25063553 | A model of the hormonal control of hepatic metabolism using insulin and glucagon signaling | Metabolic | Metazoan | 6.431252 |
| 26163548 | A signaling pathway model describing the interaction between a drug and cellular outcomes in multiple myeloma cells | Signaling | Metazoan | 5.942066 |
| 29206223 | A model to predict different in-silico knockouts that prevent key SASP-mediators, IL-6 and IL-8, from getting activated upon DNA damage | Biochemical | Metazoan | 5.414375 |
| 26446703 | A model of colitis-associated colon cancer involving P53, MDM2, and AKT | Disease | Metazoan | 4.918872 |
| 19144179 | A model of glucose regulation in yeast | Metabolic | Primitive | 4.804903 |
| 30530226 | A model of the crosstalk between the TGF-β, p38 MAPK and cell-cycle checkpoint pathways which qualitatively describes this dual behavior of TGF-β | Signaling | Metazoan | 4.597779 |
| 21563979 | A model of glucose regulation via the Lac operon | Metabolic | Primitive | 3.447058 |
| 23049686 | A model of cell-cycle in budding yeast | Regulatory | Primitive | 3.288717 |
| 23868318 | A model of receptor signaling pathways, including EGFR, G-protein-coupled receptor, integrin, and stress pathways | Regulatory | Metazoan | 2.027278 |
| 17010384 | A model used to simulate a molecular pathway between two neurotransmitter receptors: dopamine and glutamate receptors | Signaling | Metazoan | 0.686516 |
| 23171249 | A regulatory model of the TOL multi-protein complex system found on the cell membrane of gram negative bacteria | Metabolic | Metazoan | 0.508221 |
| 23868318 | A model of interaction between nine key signaling pathways in drosophila | Regulatory | Metazoan | 0.074953 |
| 16464248 | A model of a typical signaling pathway where mass and signal flow occur simultaneously | Signaling | Metazoan | 0 |
| 18194572 | A logical model of HGF and H. pylori induced c-Met signal transduction | Disease | Metazoan | 0 |
| 27594840 | A model linking the hematopoietic stem progenitor cells and mesenchymal stromal cells in the bone marrow | Disease | Metazoan | -0.19119 |
| 23520449 | A model of body segmentation in drosophila melanogaster | Biochemical | Metazoan | -0.30788 |
| 23764028 | A model of the effect of TGF-b1 on misquitos with the parasite Plasmodium falciparum | Disease | Metazoan | -0.71147 |
| 21639591 | A model for phospholipase C-coupled calcium signaling pathways | Signaling | Metazoan | -0.76057 |
| 18433497 | A model of cellular immune response integrating toll-like receptor, interferon, NF-kappaB and apoptotic pathways | Signaling | Metazoan | -1.00938 |
| 19025648 | A model of physiological regulation in cholesterol biosynthesis | Metabolic | Metazoan | -1.24445 |
| 30546316 | A model for the generegulation driving macrophage polarization to the M1, M2a, M2b, and M2c phenotypes | Signaling | Metazoan | -2.01878 |
| 23469179 | A model of cancer network regulated by miR-17-92 cluster | Disease | Cancer | -2.71715 |
| 19118495 | A regulatory model of the cell-cycle involving the transmembrane tyrosine kinase ERBB2 | Disease | Metazoan | -3.46556 |
| 23134720 | A model linking the oxidative stress pathway to apoptosis | Biochemical | Metazoan | -3.86379 |
| 26346668 | A model is used to map environmental and flow induced signals to endothelial cell phenotype (proliferation,migration, apoptosis, and lumen formation) | Disease | Cancer | -4.09752 |
| 32870080 | A model of the lac operon with feedback | Regulatory | Primitive | -4.53977 |
| 26090929 | An integrated regulatory and signaling pathway model of the fate of the CD4+ T immune cells | Biochemical | Metazoan | -6.53616 |
| 20659480 | ABA-induced closure model (ABA signal transduction network) | Signaling | Plants | -9.87253 |
| 25980672 | A model of the melanogenesis signaling network | Signaling | Metazoan | -9.90163 |
| 21968890 | A model of signaling pathway involving the pro-inflammatory cytokines interleukin 1 (IL-1) | Signaling | Metazoan | -13.6196 |
| 31048917 | A model of *Drosophila melanogaster* | Regulatory | Metazoan | -13.6436 |
| 23868318 | A model of interaction between nine key signaling pathways in *drosophila* | Regulatory | Metazoan | -13.9899 |
| 21968890 | A model of signaling pathway involving the pro-inflammatory cytokines interleukin 6 (IL-6) | Signaling | Metazoan | -14.0946 |
| 19622164 | A model of the external signaling molecule TNFa | Disease | Metazoan | -17.2082 |
| 19622164 | A model of the external signaling molecule TNFb1 | Disease | Metazoan | -17.4249 |
| 28455685 | A model of the epidermal growth factor receptor signaling of a breast epithelial cell line, MFC10A. | Disease | Cancer | -18.4117 |
| 30323768 | A model gene regulatory network (GRN) of latent proviruses in resting CD4+ Tcell to visualize the complexity of the HIV-1 gene expression | Disease | Metazoan | -20.8165 |
| 26528548 | A regulatory network model of early metasis development | Disease | Cancer | -78.6659 |
| 20862356 | A gene regulatory model of spatial expression patterns in the cerebral cortex | Regulatory | Metazoan | -92.6647 |

